# Glutathione impacts both *Batrachochytrium dendrobatidis* virulence and amphibian cellular defence in a chytridiomycosis model

**DOI:** 10.64898/2026.02.25.707882

**Authors:** Rebecca J Webb, Alexandra A Roberts, Lee Berger, Jacques Robert, Lee Skerratt

## Abstract

Glutathione has important roles in diverse infections, yet its involvement in the interaction between the deadly fungal pathogen *Batrachochytrium dendrobatidis* (*Bd*) and its amphibian hosts is still unclear. Using *in vitro* assays and a cell infection model, we examined how glutathione influences *Bd* virulence traits and cellular host disease resistance. For *Bd*, inhibition of glutathione reductase rapidly killed zoospores, indicating that glutathione is essential for this pathogen. In addition, exposure to exogenous glutathione promoted the potential for virulence through accelerated and increased zoospore release. In host amphibian cells, *Bd* infection decreased intracellular glutathione content and increased reactive oxygen species, suggesting that chytridiomycosis pathogenesis may involve oxidative stress. Depletion of host glutathione before exposure to *Bd* increased infection severity and *Bd* growth, whereas amphibian cells with slightly elevated glutathione levels were partially protected against *Bd*. However, manipulation of host glutathione levels after the establishment of *Bd* infection did not impact its intracellular growth, implying that the host glutathione-mediated resistance only occurs during the initial *Bd* invasion process. Importantly, this effect of glutathione on host resistance is not a general response to pathogens, as it was not observed in cells exposed to viral pathogen FV3. As glutathione increased both infectious zoospore production and host resistance to zoospore infection, our study suggests that this antioxidant may play an important role in the host/pathogen interaction during chytridiomycosis. Thus, environmental conditions and therapeutic approaches that affect glutathione systems in the host and/or pathogen have the potential to alter chytridiomycosis dynamics and should be further explored.

## 1. Introduction

Glutathione can influence host-pathogen interactions in multiple ways. The ubiquitous presence of glutathione in eukaryotic cells can serve as a signalling “virulence switch” for some pathogenic bacteria providing a cue that they have reached a favourable host environment (1, 2). This has not been described in fungal pathogens however, and the roles of extracellular or exogenous glutathione for modulation of virulence has not been thoroughly explored (2). Endogenous glutathione is known to be important for fungal development, and changes in the intracellular glutathione concentration or ratio can trigger processes such as germination, conidiation and dimorphic transition (3-5). Of particular importance is the maintenance of redox state achieved via glutathione reductase (GR) mediated regulation of the ratio of reduced (GSH) and oxidised glutathione (GSSG). Without GR activity, fungi can display reduced tolerance of stress, reduced virulence or complete inability to grow (6, 7). Understanding the role of fungal glutathione systems in disease can therefore inform new antifungal strategies (8).

From the host point of view, the glutathione system is also required as a means to fight infection. Host-produced glutathione is associated with resistance to viral and bacterial diseases (9, 10). Glutathione is also involved in defence against fungal infections, as GR deficient mice displayed higher *Candida* infections (11), and in cress plants, glutathione is required for resistance to the fungal pathogen *Colletotrichum gloeosporioides* (12).

*Batrachochytrium dendrobatidis* (*Bd*) is a fungal pathogen responsible for the disease chytridiomycosis, which has caused dramatic declines and extinctions of hundreds of amphibian species worldwide (13). The loss of amphibians from the ecosystem has caused trophic cascade biodiversity loss (14) and could impact human health via reduced predation of malaria transmitting mosquitos (15). Strategies to reduce the impact of chytridiomycosis, either by increasing host resistance, or decreasing pathogen virulence, are therefore urgently needed (16). During chytridiomycosis, the fungal zoosporangia grow within amphibian epidermal cells and release infective zoospores. Mucin on the amphibian skin surface likely stimulates the waterborne zoospores to recognise the host and encyst (17). Once inside host tissue, many putative virulence genes are upregulated *in vivo* (18, 19). However, the factors that trigger these transcriptional changes during intracellular infection have not yet been resolved. We previously found that exposure to exogenous glutathione resulted in a dramatic increase in zoospore production, even in zoosporangia depleted of their endogenous glutathione supplies (20). Amphibian skin cells contain glutathione (21), however the role that host-derived glutathione plays in chytridiomycosis susceptibility and disease progression is unknown. If *Bd* responds to host cytosolic glutathione via accelerated and increased production of infective zoospores, this may trigger increased pathogen virulence. Conversely, amphibian glutathione may increase resistance to fungal infection, as seen in other disease systems (22). A recent study of newly metamorphosed frogs observed lower levels of glutathione in the skin and liver of *Bd* infected animals (23), suggesting a role in the disease process. However, direct evidence of the impact of host glutathione availability on chytridiomycosis is lacking and requires experimental manipulation of host glutathione and quantification of disease outcomes.

The objective of the present work was to investigate the effect of host- and pathogen-derived glutathione on *Bd* growth and infection. Since GR can influence the growth and virulence of other fungi, we used a potent GR inhibitor to disrupt GSSG-GSH recycling and monitored the effect on growth in culture to assess whether maintenance of the reduced:oxidised glutathione ratio is essential in *Bd*. Since exposure to exogenous glutathione can trigger virulence traits in bacterial pathogens, we exposed zoosporangia to reduced and oxidised glutathione and monitored for changes in zoospore production to understand the impact of glutathione on the *Bd* life cycle and virulence potential. To assess if this response was specifically due to glutathione, we compared these results to an alternative reducing agent (ß-mercaptoethanol, BME) and an alternative extracellular amphibian skin component (mucin). A host cell infection model was used to explore how host-derived glutathione impacts fungal virulence. We monitored the change in host glutathione content during *Bd* infection and assessed fungal infection burden when host glutathione levels were manipulated. To determine whether the effect of glutathione levels on *Bd* infection was a generalised response to pathogens, we repeated the glutathione manipulation experiments with an amphibian viral pathogen. As glutathione is often utilised in response to oxidative stress, we assessed whether reactive oxygen species (ROS) were generated during *Bd* infection and explored whether ROS originated from the host or pathogen cells.

## 2. Methods

### 2.1. Chemicals

All chemicals were purchased from Merck. GSH, GSSG, BME, mucin, cysteine and buthionine sulphoximine (BSO) stock solutions were freshly prepared in MilliQ water and filter sterilised. The GR inhibitor 2-Acetylamino-3-[4-(2-acetylamino-2-carboxy-ethylsulfanylthiocarbonylamino)phenylthiocarbamoylsulfanyl]propionic acid (2-AAPA) was prepared in DMSO, and then further diluted in MilliQ water and filter sterilised to produce a 1 mM stock solution. The non-specific ROS probe 2′,7′-Dichlorodihydrofluorescein diacetate (DCFH-DA) was prepared as a 100 mM stock in DMSO and stored as frozen aliquots. The glutathione probe Monochlorobimane (mBCI) was prepared as a 50 mM stock in DMSO and stored as frozen aliquots.

### *2*.*2. Bd* culture

Three different *Bd* isolates were used in these experiments to ensure that the impact of glutathione on virulence was not specific to a single isolate. The *Bd* isolates (#46 Waste point- Litoria verreauxii #5-2013-LB, RW, #109 Melbourne-Lissotriton vulgaris-2023-LB, and #87 NarielValley-Litoria.spenceri-2020-LB) were isolated from naturally infected amphibians in Australia, and all belong to the virulent Global Pandemic Lineage (GPL). Isolates were maintained in in tryptone, gelatine hydrolysate and lactose media (TGhL) following standard laboratory protocols (24). To enable real-time visualisation of the fungal cells as they infected and grew inside frog cells, *Bd* isolate #87 was transformed with a fluorescent reporter (25) and renamed T#5. Briefly, zoospores were electroporated to deliver a plasmid conferring tdTomato fluorescence and hygromycin resistance, and kept under hygromycin selection to produce a red fluorescing *Bd* isolate.

### 2.3. Effect of disruption of endogenous glutathione on zoospores

The potent GR inhibitor (2-AAPA) was used to disrupt the endogenous glutathione pool by preventing the conversion of GSSG to GSH. A suspension of zoospores (#87) was collected from actively growing flasks, filtered through a 10 µM syringe filter (Millipore), centrifuged, then added to 96 well plates at 5× 10^4^ zoospores in TGhL per well with 10 µM 2-AAPA, or 0.05% DMSO control. A subset of wells also contained either 1.5 mM GSH or 1.5 mM GSSG. After 30 mins exposure, zoospore’s survival was determined by observing movement under a microscope (a total loss of motility was considered as indicative of zoospore death). After 3 hr, the exposure solution was removed and replaced with 100 µL of fresh TGhL, and incubated at 20°C. Relative growth of exposed zoospores was measured after 48 hr using a methylene blue growth assay to determine the fungal biomass (26). Two-way ANOVA and Tukey’s post hoc tests were used to investigate the interaction between treatment with 2-AAPA and supplementation with either GSH or GSSG.

### 2.4. Effect of exogenous glutathione on *Bd* growth

GSH, GSSG, BME, and mucin were used to determine if they elicit a potential virulence switch in *Bd*. Standard minimal inhibitory concentrations (MIC) assays (27) were conducted to determine sensitivity to each chemical, and appropriate concentrations chosen for the remaining experiments. Zoospores (#46) were plated and left to encyst for 18 h as described above. Excess media was removed and replaced with 100 µL TGhL containing either GSH and GSSG (0.5 mM-2 mM), BME (0.1 mM-2 mM) or mucin (0.25 mg/mL-2.5 mg/mL) and incubated at 20°C. After an additional 48 h incubation, an aliquot (10 µL) from three replicate wells was used to quantify zoospore production using a haemocytometer. Additional time course experiments were conducted to investigate the timing and duration of zoospore production after exposure to GSH and GSSG. Zoospores were incubated for 48 h before exposure to either 2 mM GSH or 2 mM GSSG, and zoospore production calculated by counting the next generation of motile zoospores at various time points (24 h – 90 h) post glutathione exposure. One way ANOVA and Dunnett’s post-hoc test was used to determine differences in logged zoospore count at each time point. The viability of zoospores produced following GSH exposure was assessed by transferring a 20 µL aliquot to a new well with 80 µL TGhL, allowing zoospores to encyst, then replacing the TGhL to remove residual traces of glutathione.

Relative growth was measured via methylene blue growth assay after 48 h incubation at 20°C. One way ANOVA and Dunnett’s post-hoc test was used to determine differences in growth in zoospores obtained from GSH treated zoosporangia compared to those from the untreated control.

### 2.5. Effect of *Bd* infection on host glutathione

To explore the role of glutathione for the host response to *Bd* infection, we took advantage of an established chytridiomycosis cell infection model utilising commercially available immortalized amphibian “A6” cells (derived from *Xenopus laevis* kidney) (25, 28, 29). Total glutathione was measured in A6 cells after exposure to *Bd* at two multiplicity of infection (MOI) rates. A6 cells were maintained in 75 % NCTC-109 media supplemented with 10% FCS and 2 mM glutamine at 27°C and 5% CO_2_ following standard protocols. At the beginning of the experiment, A6 cells were detached with trypsin, and added to 96 well plates at 9 × 10^3^ cells per well. After 18 h incubation at 27°C, excess media was removed from the wells and replaced with 100 µL of solution C containing either 9 × 10^3^ (1 MOI) or 2.7 × 10^4^ (3 MOI) zoospores per well for 1 hr, after which the inoculation solution as removed and replaced with 100 µL solution A (25, 28). The total glutathione concentration of host cells was measured at 4 h and 24 h after infection using a commercial kit (GSH-Glo™ Glutathione Assay-Promega) following the manufacturer’s directions. Two zoospore only wells were included to estimate the glutathione contribution from *Bd* cells, and this was subtracted from the infected wells. One Way ANOVA and Dunnett’s post hoc tests were used to determine whether host cell glutathione changed in response to *Bd* infection. A separate experiment was conducted to confirm the effect of *Bd* infection using (mBCI) a glutathione fluorescent probe which emits blue fluorescence when conjugated with glutathione (30). A6 cells were infected with 3 MOI zoospores as above and incubated at 20°C. After 72 hr, excess media was removed, and cells were washed with “amphibian strength PBS” (APBS), and then stained with 50 µM mBCI in 70% L15 media for 1 hour (31).

The staining solution was removed, cells washed with APBS and each well refilled with 100 µL solution A. Glutathione was visualised under an inverted fluorescent microscope (EVOS 5000) using a DAPI filter.

### 2.6. Effect of *Bd* infection on ROS

A similar experimental design was employed to investigate the production of reactive oxygen species (ROS) during infection. Initially, cells were infected with four different zoospore (#87) loads, 0 MOI, 0.5 MOI, 1 MOI and 2 MOI, and incubated at 20°C for 4 days. ROS was visualised with using DCFH-DA, a non-specific redox probe that emits green fluorescence in the presence of any type of ROS. Excess media was removed and replaced with 70% L15 media containing 10 µM DCFH-DA and incubated at 20°C for 30 min (32), after which the solution was removed, replaced with 10 μg/ml Calcofluor White (CFW) to stain *Bd* for 5 mins (28), rinsed and replaced with 70% L15 media. Cells were imaged under GFP fluorescence to semi-quantitatively assess ROS levels. To investigate whether ROS was host or pathogen generated, a separate experiment was conducted in which cells were infected with 1 MOI zoospores (T#5), and either: 1) supplemented with 2.5 mM cysteine on day 3 for 8 hr, 2) incubated at 27°C for 8 hr, 3) supplemented with 1.5 mM cysteine and incubated at 30°C for 6 hr. ROS staining was conducted as above to determine whether ROS levels decreased under these conditions. To examine the interaction between glutathione levels and ROS, infected A6 cells were first stained with 50 µM mBCI in 70% L15 media for 1 hour, then stained with 10 µM DCFH-DA for 30 mins.

### 2.7. Manipulation of host cell glutathione

The feasibility of manipulating host glutathione was determined using A6 cells. A6 cells were maintained in 75 % NCTC-109 supplemented with 10% FCS and 2 mM glutamine at 27°C and 5% CO2 as above. A6 cells were detached with trypsin, added to 96 well plates at 6 x10^3^ cells per well and incubated at 27°C overnight. To deplete or increase cellular glutathione, A6 cells were exposed to 10 mM BSO (33) (Shirriff and Heikkila, 201 or 2.5 mM cysteine (34, 35) for 18 h respectively. Total cellular glutathione was measured using a commercial kit (GSH-Glo™ Glutathione Assay-Promega) following the manufacturer’s directions.

One way ANOVA analysis with Dunnetts post-hoc tests were used to determine if total cellular glutathione responded to treatment. A separate experiment was conducted to confirm glutathione manipulation was successful using mBCI to stain glutathione as described above. A6 cell viability upon BSO and cysteine treatment was determined using MTT assay. A6 cells were added to 96 well plates at 2.5 x10^4^ cells per well, incubated at 27°C overnight, then exposed to 10 mM BSO or 2.5 mM cysteine for 18 hours, before determining cell viability using CyQUANT™ MTT assay following the manufacturer’s instructions.

### 2.8. Effect of host derived glutathione on *Bd* growth and host cell health

To explore the effect of host glutathione levels on *Bd* infection, A6 cells were manipulated as above to increase or decrease glutathione levels before and after exposure to *Bd* zoospores. Actively growing A6 cells were detached with trypsin, added to 96 well plates at 2.5 x10^4^ cells per well and incubated with 10 mM BSO or 2.5 mM cysteine to alter glutathione levels. After 18 h incubation at 27°C, excess media was removed from the A6 wells and replaced with 100 µL of solution C containing 5 × 10^4^ T#5 zoospores (2 MOI). After 2 hours incubation at 20°C, the zoospore solution was removed and replaced with 100 µL of solution A, along with either 10 mM BSO or 0.25 mM cysteine and incubated at 20°C. Cysteine concentrations were reduced after *Bd* introduction as MIC experiments in TGhL indicated that concentrations above 1 mM affect *Bd* growth, whereas BSO doesn’t inhibit *Bd* growth until 30 mM (20).

After 24 h, the media was replaced with solution A, and the cells left to grow for an additional 6 d. To understand the effect of host cell glutathione on existing infections, additional experiments were conducted in which A6 cells were supplemented with 2.5 mM cysteine at 24 hr post infection. *Bd* growth was assessed by taking 3 randomly chosen images at 4x magnification fields of view (FOV) per well under red fluorescence, and calculating the total area occupied by the red fluorescing *Bd* cells using image J relative to the control average (36). If wild type *Bd* was used to infect the cells, *Bd* growth was measured using calcofluor white (28). *Bd* zoospore production was measured by counting the number of motile zoospores present at 3 randomly chosen FOV per well and calculated relative to the control average. A6 cell health was determined by staining with 50 µL sterile filtered 0.5 µg/mL DAPI (Invitrogen) for 5 min, after which the staining solution was removed and replaced with 100 µL solution A. Uninfected controls were included to check for damage from cysteine or BSO treatment alone. Host cell damage was measured by imaging under blue fluorescence and the number of DAPI positive cells calculated using Image J, relative to the untreated control average, using the uninfected treatment as a blank. One way ANOVA analysis with Dunnett’s post-hoc tests were used to determine if *Bd* growth, zoospore production or host cell damage differed from untreated control cells.

### 2.9. Effect of host derived glutathione on disease susceptibility of host cells to Frog Virus 3

To explore whether changes in host cell glutathione cause a generalised effect on disease susceptibility, the above experiment was repeated with a ranavirus as an alternative amphibian pathogen. After 18 h BSO and cysteine incubation, excess media was removed, replaced with 100 µL A6 media containing 0.6 MOI frog virus 3 (FV3) (37), and incubated at 30°C for 1 hour. After viral exposure, excess media was removed and replaced with fresh A6 media containing 10 mM BSO or 2 mM cysteine and incubated at 27°C for 72 h. At 56 h post infection A6 cell survival was measured using a CyQUANT™ MTT Cell Proliferation Assay Kit (Thermo) following the manufacturer’s instructions and calculated relative to the FV3 only control.

### 2.10. Statistical analysis

All data were obtained from *in vitro* experiments in 48 or 96 well plates, with each individual well considered a pseudo-biological replicate. Multiple measurements (eg; counts or images) from the same well were treated as technical replicates and averaged. Data were analysed using Graphpad Prism 10.2.3. Count data (zoospore counts and damaged cell counts) were logged transformed. One way, two way or repeated ANOVA and appropriate post-hoc tests were used to identify significant differences, with a significance level set at p < 0.05. All test assumptions of normality were met. Graphical data is expressed as mean ± standard deviation.

## 3. Results

### 3.1. Decreased GSH levels via inhibition of glutathione reductase is acutely toxic to *Bd*

We have previously found that chronic inhibition of endogenous glutathione synthesis prevents *Bd* growth and development *in vitro* (20). Here we found that inhibition of the enzyme responsible for endogenous glutathione recycling (GR) is acutely toxic (Figure 1). There was a significant interaction between 2-AAPA treatment and the type of glutathione supplementation (ANOVA F_2,12_ = 104.7, p=<0.0001). Exposure to 10 µM 2-AAPA caused rapid toxicity to zoospores, resulting in a rapid loss of motility, and significantly less growth than the untreated control (p= <0.0001). In 2-AAPA treated wells, supplementation with 1.5 mM GSH provided a partial rescue effect, with growth significantly higher than the non-supplemented 2-AAPA treated control (p=<0.0001), whereas 1.5 mM GSSG did not reverse the effects of 2-AAPA exposure and growth was not significantly different to the non-supplemented 2-AAPA treated control (p=>0.9999).

**Figure 1:**
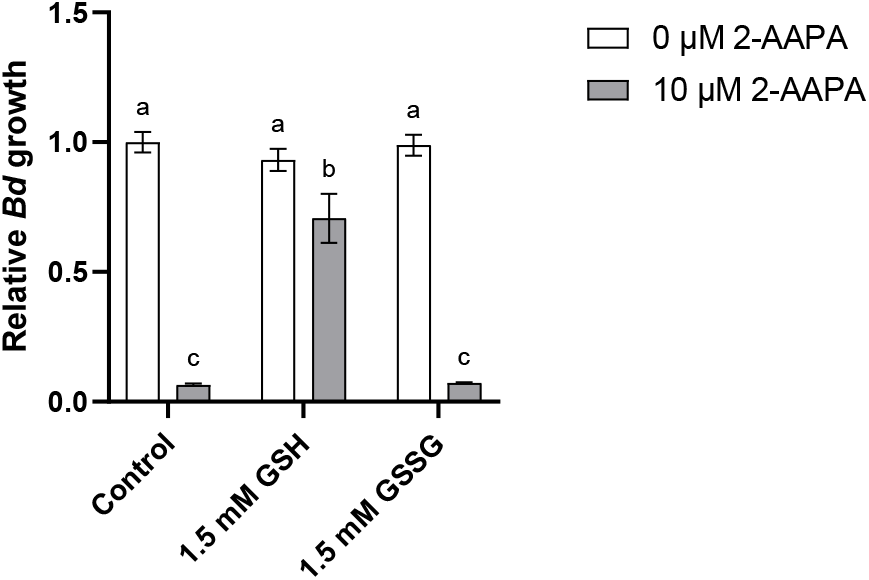
The toxic effects of 2-AAPA can be rescued by supplementation with GSH, but not GSSG. 2-AAPA is a potent glutathione reductase inhibitor, preventing the conversion of endogenous oxidised glutathione (GSSG) back to the reduced form (GSH). *Bd* zoospores were incubated with 10 µM AAPA with or without GSH or GSSH. Bars show mean growth and SD of 3 replicate wells measured 24 hours after 2-AAPA exposure using a methylene blue growth assay. Bars that do not share a common superscript letter differ significantly according to a two-way analysis of variance (ANOVA) for independent samples, followed by Tukey’s post-hoc test (p < 0.05), n=18

### 3.2. Exogenous glutathione triggers increased zoospore production in *Bd*

To investigate potential virulence triggers, young zoosporangia (20 h) were exposed to GSG, GSSG, BME or mucin and monitored for zoospore release at various timepoints after exposure. Both oxidised and reduced glutathione triggered an increase in viable zoospore production relative to the untreated control (Figure A1). An increase in zoospore production was not observed in zoosporangia exposed to BME (another thiol reducing agent), or mucin (a component of the amphibian skin mucosome). To substantiate this finding, mature zoosporangia (48 h) were exposed to different concentrations of GSH or GSSG and zoospore release monitored over time (Figure 2). Glutathione appeared to act as a catalyst that accelerated zoospore release by at least 12 h compared to the control, and this increased zoospore production was sustained during the growth cycle (Figure 2). There were significantly more zoospores released by zoosporangia exposed to GSH or GSSG at 24 hr (p=0.0155, 0.0342 respectively), 48 h (p=0.0310, 0.0344 respectively), 66 h (p=0.0063,0.0279 respectively). By 90 hr post exposure, zoospore release was not significantly different to that of the control.

**Figure 2:**
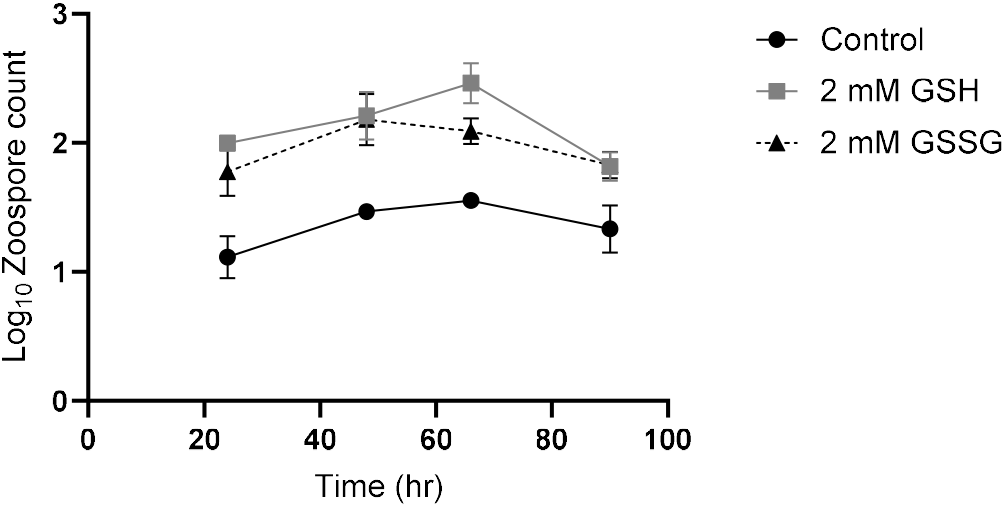
Increased zoospore production is sustained after exogenous glutathione exposure. Zoospore production of mature (2 d old) *Bd* zoosporangia exposed to 2 mM reduced (GSH) or oxidised (GSSG) glutathione compared to an untreated control over one lifecycle. Mean logged zoospore counts estimated via haemocytometer from two replicate wells per treatment per timepoint and SD, n=24.

### *3*.*3. Bd* infection depletes host glutathione stores and increases ROS

An *in vitro* cell infection model (25, 28) was used to further investigate the role of glutathione during chytridiomycosis. Infected amphibian (A6) cells had lower total glutathione levels compared with uninfected controls (Figure 3A). Using a luminescent glutathione assay, we found exposure to low zoospore burdens significantly decreased host glutathione content by 36% after 24 h (p=0.0058). At high zoospore burdens, host glutathione content was significantly decreased by 38% after 4h (p=0.0038) and 41% after 24 h (p=0.0021). Visualisation of glutathione content using 50 µM mBCI produced similar results, with infected cells displaying lower staining intensity (Figure 3B). Infection also increased amphibian cell ROS in a dose dependent manner (Figure 3C). ROS appeared to be closely associated with *Bd* infection, as DCFH-DA staining was concentrated in cells containing large intracellular zoosporangia (Figure 3D, A2). Staining shows *Bd* zoosporangia contained more glutathione and less ROS compared to the A6 cells (Figure 3D).

**Figure 3:**
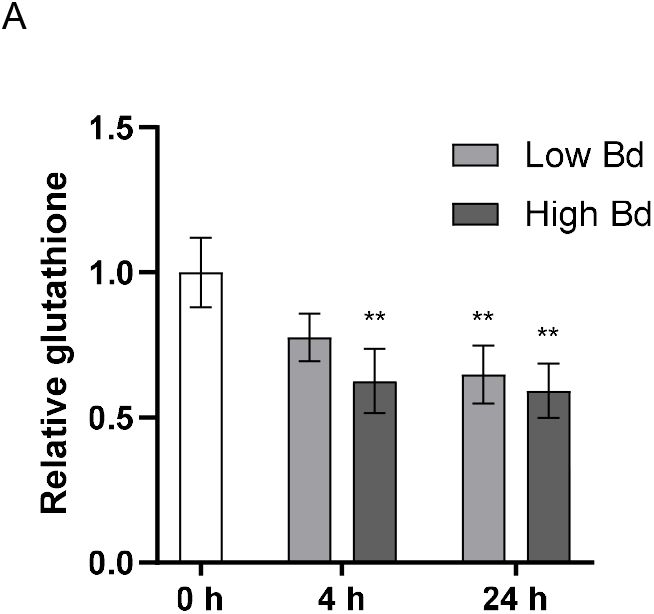
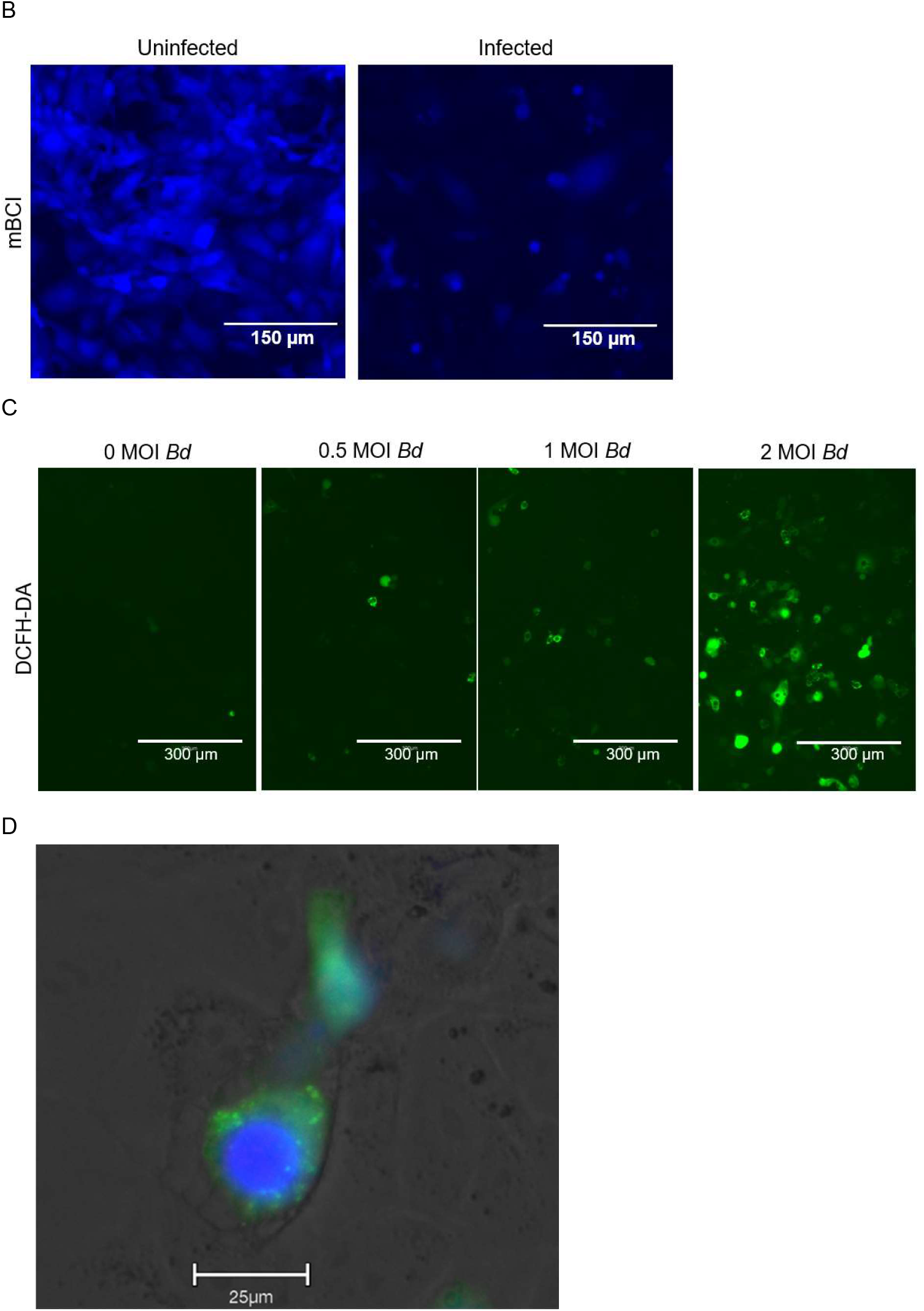
*Bd* infection causes a decrease in host cell glutathione content and increase in ROS. **(A)** The total glutathione abundance of immortalised amphibian cells (A6) was measured with a luminescent GSH assay (Promega) before and during infection with either a low (1 MOI, multiplicity of infection) or high (3 MOI) concentration of zoospores. Mean total glutathione levels and SD of 3 replicate wells per treatment and timepoint are displayed. One way ANOVA was used to determine significant differences from the 0 h control time point indicated by asterisks (** P ≤ 0.01), n=15. (**B**) Glutathione content of uninfected and infected A6 cells was visualised using 50 µM Monochlorobimane (mBCI). Representative image at 20x magnification. (**C)** The ROS content of uninfected A6 cells compared to cells infected with 0.5 MOI, 1 MOI or 2 MOI zoospores was visualised using 2′,7′-Dichlorodihydrofluorescein diacetate (DCFH-DA), representative images at 10x magnification. **(D)** Combining both stains indicates that *Bd* zoosporangia (arrow) stain strongly for glutathione (blue) compared to the host cells which stain strongly for ROS (green).

To investigate whether the ROS is generated by the host or the pathogen, we inhibited *Bd* growth by increasing the temperature beyond the thermal optimum of *Bd*, and/or supplemented the cells with cysteine (a glutathione precursor) with the hypothesis that if ROS was pathogen generated it should decrease under these conditions. None of these treatments resulted in decreased intracellular ROS (Figure A2). Together these results suggest that the observed ROS is likely generated by the host cells rather than *Bd*.

### 3.4. Host glutathione availability affects susceptibility to *Bd* but not Frog Virus 3 (FV3)

We achieved manipulation of glutathione levels in A6 cells using BSO and cysteine. Total glutathione concentrations decreased by approximately 70% after 18 hr incubation with 10 mM BSO and increased by approximately 25% after incubation with 2.5 mM cysteine, and these effects were not due to impacts on A6 viability (Figure A3). To investigate the effect of host cell glutathione concentration on disease parameters, we infected cells that had been pre-incubated with either BSO or cysteine and compared *Bd* growth to that within untreated cells. To distinguish whether host cell glutathione concentration is important during initial infection or subsequent *Bd* growth, an additional treatment group included A6 cells that were only incubated with BSO immediately after infection. The availability of glutathione in the host cells impacted the *Bd* load (zoosporangia and zoospores) as well as host cell damage (Figure 4A,C). Host cells with elevated glutathione had significantly less zoospores present compared to control cells (p=0.0194). In contrast, host cells with depleted glutathione stores prior to infection supported significantly more *Bd* growth (p=0.0.125) and zoospore production (p=0.0018) compared to the control. Host cells that experienced glutathione depletion only after *Bd* zoospore encystation did not differ from the untreated control (Figure 4A,C). To investigate the effect of glutathione on host cell health during infection, we assessed host cell damage using DAPI, which stains the nuclei of cells with compromised membranes (36). The relative number of DAPI stained cells followed a similar pattern to *Bd* growth (Figure 4A,C). Manipulation of host cell glutathione with BSO or cysteine did not change A6 cell survival when exposed to an alternative amphibian pathogen, FV3 (Figure 4B). To investigate the effect of glutathione on existing intracellular infections, we supplemented A6 cells with 2.5mM cysteine 24 hr after infection. Cysteine supplementation restored glutathione levels in infected cells (Figure 4D) but had no significant effect *Bd* growth (Fig A4).

**Figure 4:**
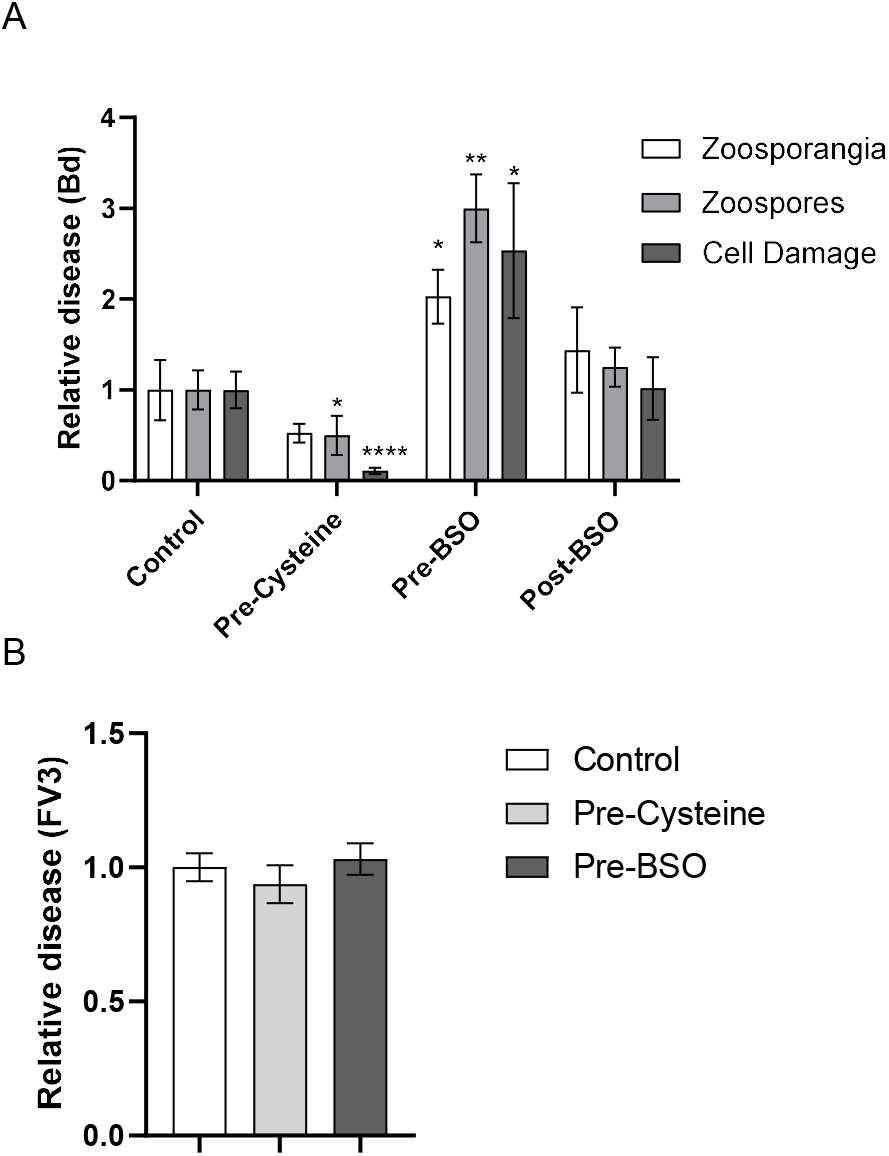
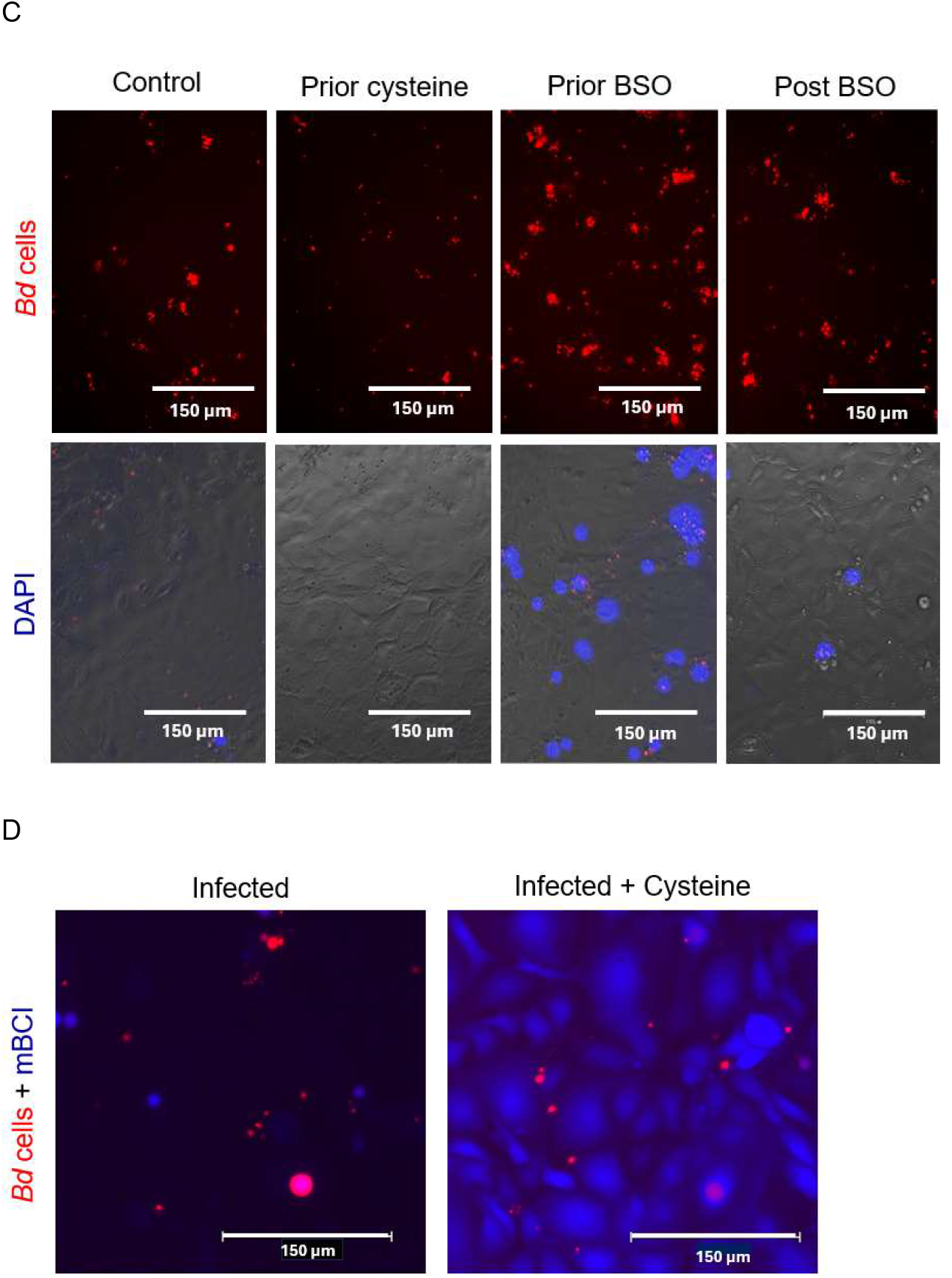
Glutathione availability in host cells affects chytridiomycosis. Immortalised amphibian cells (A6) with varying glutathione concentrations were exposed to *Bd* zoospores or Frog Virus 3 (FV3) to assess differences in host cell damage and *Bd* infection burden as measures of disease. Glutathione content of A6 cells was increased prior to *Bd* infection via supplementation with 2.5mM cysteine, and depleted either prior or after infection via addition of 10 mM BSO. **(A)** *Bd* growth was measured as the red fluorescence surface area (zoosporangia) or zoospore count (zoospores) at 7 d. A6 cell damage was assessed by the number of DAPI positive cells at 3 d. Mean values from three replicate wells and SD are displayed, One way ANOVA and Dunnett’s post-hoc tests was used to determine significant differences from the control (* p= ≤0.05, ** p ≤ 0.01, **** p≤0.0001), n=12. **(B)** A similar experiment in which A6 cells were exposed to 2.5 mM cysteine or 10 mM BSO before challenge with FV3 ranavirus. Relative disease was calculated by MTT assay at 3 d. Mean cell activity from three replicate wells and SD are displayed, n=12. **(C)** Representative images of *Bd* infection at 20x magnification show *Bd* growth (red) after 7 d, or DAPI positive A6 cells (blue) after 72 h. Cells received either no treatment (control), 2.5 mM cysteine prior to infection (Prior cysteine), 10 mM BSO prior and after infection (Prior BSO), or 10 mM BSO only after infection (Post BSO). **(D)** Supplementation with cysteine 24 hrs after *Bd* infection restored depleted glutathione levels, as indicated with mBCI staining, but did not reduce *Bd* growth. Representative images at 20x show glutathione (blue) and *Bd* cells (red) 4 days after infection.

## 4. Discussion

Our study provides compelling evidence that glutathione has a fundamental role in the pathogenesis of chytridiomycosis, impacting traits of both the pathogen and the host. We also show for the first time that ROS increases within infected cells *in vitro*.

Exposing *Bd* to glutathione *in vitro* triggered a dramatic and sustained increase in zoospore production, indicating that glutathione, ubiquitous within host cells, may act as a cue to signal a favourable host environment. We also show that the maintenance of redox balance via glutathione recycling is an essential process in *Bd*, as inhibition of GR activity is acutely toxic. For the host, increased glutathione availability promotes resistance to *Bd* infection, possibly due to its role in protecting the host from ROS generated in response to fungal invasion.

### 4.1. Effect of glutathione on *Bd*

*In vitro, Bd* zoosporangia responded to low levels of exogenous glutathione via a dramatic and sustained increase in zoospore production. Preliminary experiments indicated that the concentration of GSH needed to trigger increased zoospore release varied slightly depending on the *Bd* isolate. The timing of glutathione exposure does not appear to be crucial, as zoosporangia responded in a similar manner when exposed at 24 or 48 h. Accelerated zoospore production has been correlated with high virulence strains in *in vivo* infection trials (38-42), thus our results suggest that glutathione may be an important for *Bd* virulence. The mechanism behind the stimulation of zoospore release by glutathione is not yet clear, but the observation that both GSH and GSSG elicit a zoospore response, whereas mucin or BME does not, implies that *Bd* is not merely responding to exposure to host tissue or a generalised redox change, as hypothesised in *B. pseudomallei* (43). As previous studies demonstrated, zoosporangia under severe endogenous glutathione depletion also increase zoospore production above baseline when exposed to exogenous glutathione (20). Thus, exogenous glutathione may be acting as a specific trigger to spur accelerated zoospore release.

Perhaps glutathione serves as a transcriptional activator of growth-associated genes, or that increased glutathione could facilitate differential glutathionylation of proteins essential for metabolism such as the glycolytic enzyme enolase (44). In human T lymphocytes, the glycolytic enzyme enolase is constitutively glutathionylated, which reduces its enzymatic activity. Peroxide stress increases glutathionylation of human enolase to reduce activity and likely protect the lymphocytes against oxidative stress by reducing metabolism (44). In *Bd*, enolase has also been shown to be constitutively glutathionylated, however, in contrast to human lymphocytes, peroxide stress reduces the proportion of glutathionylated enolase in *Bd* (45). Further studies are required to confirm whether changes in redox balance, and de-glutathionylation of enolase specifically, stimulates the glycolysis pathway and *Bd* growth.

Given the likely association of glutathione with *Bd* virulence, we sought to investigate the role of endogenous glutathione in *Bd*. Continual synthesis of endogenous glutathione is essential for normal growth and development (20), but the importance of maintaining the correct ratio of reduced:oxidised glutathione in *Bd* was unknown. To examine the importance of this process, we took advantage of the potent *GR* inhibitor 2-AAPA (46), commonly used at 25 µM (47). Inhibition of GR prevents *Bd* from converting GSSG into GSH, and from our previous studies on GSH depletion, we predicted that 2-AAPA would have some impacts on growth and metal tolerance (27). Unexpectedly, preventing glutathione recycling was acutely toxic to *Bd*. As opposed to inhibiting glutathione synthesis via BSO, which did not affect zoospores, and took at least 24 hours to impact growth of zoosporangia (20), inhibiting GR with 10 µM 2-AAPA killed zoospores within 30 min, and 50 µM 2-AAPA killed zoosporangia within a few hours. To confirm that the toxicity was due to depletion of GSH, we supplemented 2-AAPA treated zoospores with either GSH or GSSG (6). GSH, but not GSSG, reversed the effect of 2-AAPA, confirming the mechanism of toxicity as redox imbalance and the inability to recycle GSH from GSSG. To our knowledge, this is the first time that the use of 2-AAPA has been validated in chytrid fungi. This demonstrates that GSH and the maintenance of the redox balance is essential in *Bd*. The acute requirement for glutathione reductase activity in *Bd* is in contrast with other fungal species, in which GR knockouts display only reduced tolerance to oxidative stress (48) or temperature (6). Given that *Bd* cannot survive even a few hours without the ability to recycle glutathione, GSH represents an important molecule for *Bd*. As such GR is an indispensable enzyme and potential target for novel antifungal strategies such as RNAi (16).

### 4.2. Effect of glutathione on host cells

Glutathione also functions in protecting amphibian cells against *Bd* infection. In an A6 cell infection model, we saw increased fungal growth when host cells were deprived of glutathione via BSO at the time of infection. When A6 glutathione levels were increased via cysteine supplementation, there was a reduction in *Bd* and associated host cell damage. Since *Bd* growth *in vitro* is increased with additional glutathione, the opposite intracellular effect suggests the outcome of manipulating cell glutathione levels is a result of altered host resistance. This is consistent with host resistance to other fungal pathogens. We explored the precise timing of glutathione’s role in immunity by depleting glutathione in A6 cells before and after infection with *Bd*. Indicators of disease increased in A6 cells depleted prior to – but not after – zoospore exposure, indicating that host cell glutathione is important during initial zoospore infection. We note that the effect of host cell glutathione was not observed when A6 cells were exposed to a viral pathogen, suggesting that this is not a generalised disease response, but may be specific to chytridiomycosis. As glutathione levels were lower in *Bd*-infected A6 cells, this suggests host glutathione is being utilised for immune defence. Alternatively, *Bd* may be modulating the host glutathione to its own advantage as occurs during *Cryptococcus* infections (49).

To further investigate glutathione relevance for host resistance to *Bd* infection, we monitored cellular ROS, and found it increased in infected A6 cells. The response correlated with zoospore dose. However it was not clear whether ROS was produced by *Bd* as a virulence mechanism, as seen in many phytopathogenic fungi (50, 51), or alternatively produced by the host as a response to infection (52), or associated membrane damage (22). To help resolve this, we decreased *Bd* metabolic activity by increasing the temperature outside of its thermal optimum. In addition, we supplemented the host cells with cysteine to increase the capacity to neutralise unwanted ROS. Neither of these treatments resulted in a decrease in ROS. This preliminary evidence suggests the observed high levels of ROS are generated by the host rather than the pathogen. As DCFH-DA is a general ROS probe, the use of more specific probes or NOX-inhibitors in the future would help to further understand the response of *Bd* infection on host redox status.

Combined, these results suggest that glutathione is an important host resistance mechanism, but that also and ability to withstand host ROS, and potentially use this as a trigger for zoospore production, is important for *Bd* virulence. Indeed, the ability to scavenge host produced ROS is a virulence factor in many phytopathogenic fungi, and disruption of oxidative stress enzymes, such as superoxide dismutase (SOD), can result in lower virulence (53). This may also be the case for *Bd*, as SOD and glutathione transferase (GST), are differentially expressed between *Bd* pandemic and non-pandemic lineages (54), suggesting that managing host ROS could be related to virulence. Glutathione is utilised by *Bd* in response to oxidative stress (20), and we observed that *Bd* cells stain strongly with mBCI, indicating they contain high concentrations of glutathione. We also found that GR (needed to recycle GSSG back to GSH during ROS detoxification) is essential for zoospore viability. Together, these results point to an important role for ROS detoxification in *Bd*. Future functional genetic studies could provide direct evidence of the importance of these enzymes during the infection process and provide targets for new antifungal strategies.

Studies in other fungal pathogens have shown the potential for modulation of glutathione for biocontrol. Direct supplementation with glutathione reduced cell mortality in a *Candida albicans in vitro* infection model (22). In crops, application of chemicals (55), or genetic constructs (53) that increase GSH levels can provide strong plant protection against fungal pathogens. We found that cells supplemented with cysteine were able to restore glutathione levels and reverse the depletion caused by *Bd* infection. Future studies should investigate whether glutathione supplementation can reduce chytridiomycosis *in vivo*. Given that amphibian cellular glutathione can be impacted by environmental contaminants such as agricultural pesticides (56-58), their potential immunosuppressive effects regarding chytridiomycosis should be investigated. In contrast, exposure to environmental heavy metals can increase glutathione in amphibian tissues (59), perhaps contributing to increased survival in “metal disease refuges” (60). Data is needed on the glutathione concentration of amphibian epidermis, to compare to levels that impacted growth in the kidney cell model. Further work could also test for correlations in susceptibility to chytridiomycosis and variation in epidermal glutathione concentrations between individuals or amphibian species.

## 5. Conclusion

The current body of knowledge suggests a complex relationship between *Bd* and the glutathione system. Glutathione is important for *Bd* growth (20) and metal tolerance (27). Given the role of GR in glutathione recycling is essential for zoospore survival, targeting this process could provide an opportunity for developing a novel antifungal strategy. Glutathione also plays an important role in host disease resistance, as infection causes depletion of intracellular glutathione, while host cells with elevated glutathione levels are less susceptible to *Bd* infection. Therefore, increasing host glutathione levels may represent an opportunity to improve resistance to chytridiomycosis. We discovered that ROS increase in host cells during infection, and these ROS are likely generated by the host rather than from the pathogen itself.

Glutathione’s role as an antioxidant likely modulates this process, thus contributing to host resistance. The fact that *Bd* zoosporangia (the parasitic intracellular life stage) contain high levels of glutathione would help the pathogen to resist ROS activity from the host. Taken together, these results provide evidence that glutathione plays a complex role in the interplay between pathogen, host, and environment in the pathophysiology of chytridiomycosis. Future work is needed to evaluate whether modulation of host and/or pathogen glutathione systems represents a viable strategy for disease intervention.

## Author Contributions

**RJW:** Conceptualization (lead), Methodology (lead), Investigation (lead), Formal analysis (lead), Funding acquisition (equal), Writing-original draft (lead). **AAR** Conceptualization (supporting), Methodology (supporting), Supervision (equal), Writing-reviewing and editing (equal). **LB:** Methodology (supporting), Supervision (equal), Writing-reviewing and editing (equal), Funding acquisition (equal), Resources (equal). **JR:** Methodology (supporting), Supervision (equal), Writing-reviewing and editing (equal), Resources (equal). **LFS:** Formal analysis (supporting), Supervision (equal), Writing-reviewing and editing (equal), Funding acquisition (equal), Resources (equal).

## Acknowledgements

Thanks to Jerome Le Nours for kindly supplying the A6 cells. Thanks to Amy Aquilina, Kashmini Sumanasekera, Nazia Akram, Francisco De Jesús Andino, Laura Brannelly and Ben Cuff for laboratory support. RJW is supported by the Walter Fisher Grant for Mycology Research on behalf of the Royal Society of Queensland, and the 2024 ECR grant on behalf of the University of Melbourne, and by D25ZO-FP-303 on behalf of the Morris Animal Foundation. LB is supported by DP220101361 on behalf of the Australian Research Council, LFS is supported by FT190100462 on behalf of the Australian Research Council, JR and the Xenopus laevis Research Resource for Immunobiology are supported by R24-AI-059830 on behalf of the National Institute of Allergy and Infectious Diseases).

## Appendix

**Figure A1:**
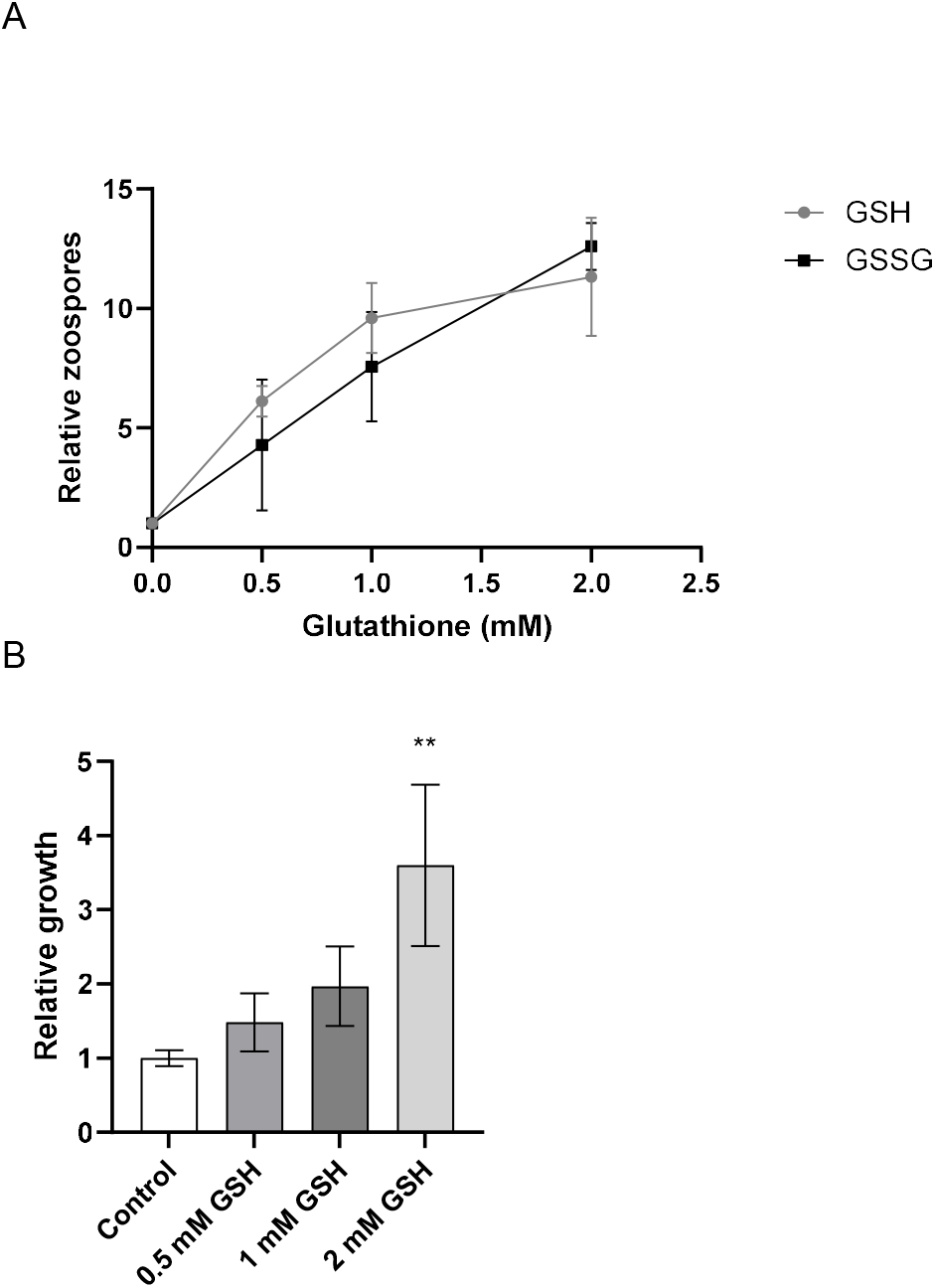
Exogenous glutathione exposure increases zoospore production. **(A)**Young (20h) zoosporangia (#46) exposed to either reduced (GSH) or oxidised (GSSG) glutathione produced more zoospores compared to control cells. Zoospores from three replicate wells per treatment were counted on a haemocytometer 48 h after glutathione exposure and calculated relative to the control. Mean zoospores per glutathione concentration and SD are displayed N=12. **(B)** To check for viability of these zoospores, an aliquot (20 µL) of these zoospores was transferred to a new plate. Zoospores released from glutathione-exposed zoosporangia encysted and grew. Bars show growth and SD of 3 replicate wells measured at 48 hours since plating using a methylene blue growth assay, relative to the control. One way ANOVA and Dunnett’s post-hoc test was used to determine differences from the untreated control, (** p ≤ 0.01), n=12.

**Figure A2:**
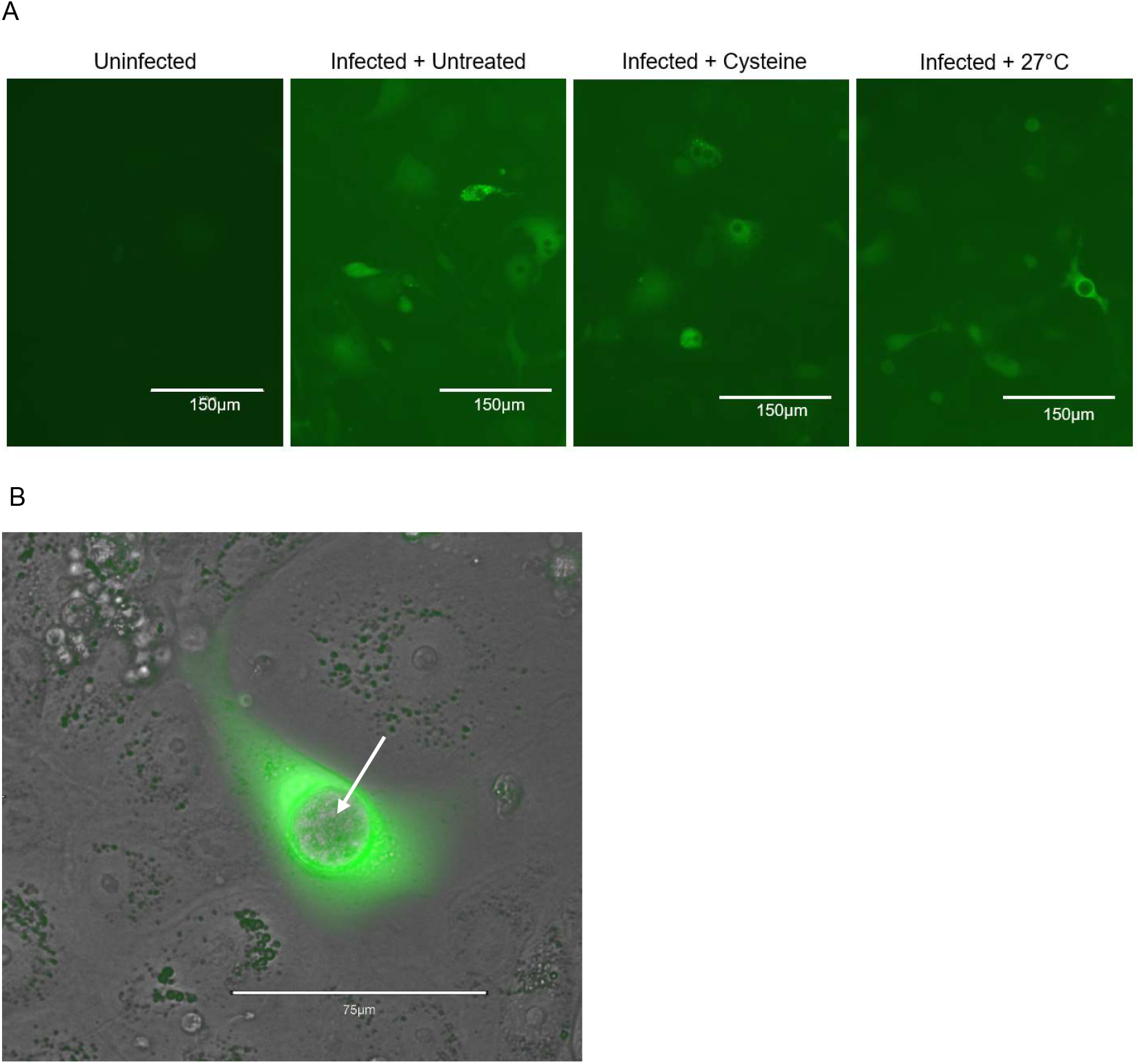
Intracellular ROS appears to be host generated. **(A)** The amount of ROS in infected A6 cells did not decrease in response to supplementation with 2.5 mM cysteine or incubation at 27°C. Uninfected cells and untreated infected cells for comparison. ROS was visualised using 2′,7′-Dichlorodihydrofluorescein diacetate (DCFH-DA), representative images at 10x magnification. **(B)** Close up image to demonstrate that ROS persists after inactivation of *Bd*. Infected A6 cells (D5 post infection) were supplemented with 1.5 mM cysteine and incubated at 30°C for 6 hours, before staining DCFH-DA for 30 mins to visualise ROS. Arrow shows intercellular Bd zoosporangia, representative image at 40x, scale bar is 75µm.

**Figure A3:**
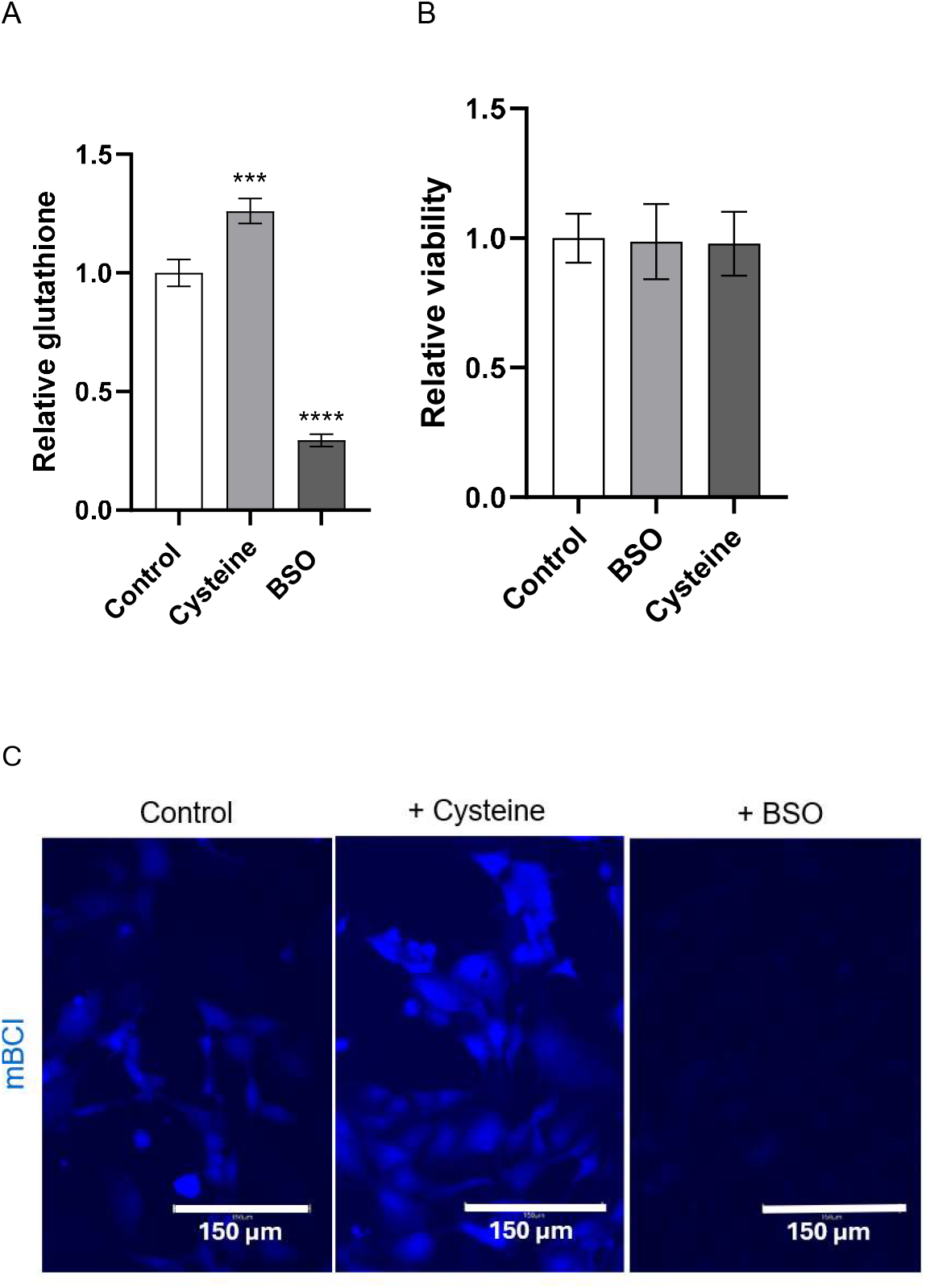
Manipulation of glutathione concentrations in A6 cells. A6 cells were incubated with 2.5 mM Cysteine (a glutathione precursor), or 10 mM Buthionine sulphoximine (BSO) (a glutathione synthesis inhibitor) for 18 h. **(A)** Bar graphs show mean and SD of total glutathione levels were measured using a luminescent quantification assay (Promega), calculated relative to the control. One way ANOVA and Dunnett’s multiple comparison test was used to determine significant differences in glutathione content from the control, (***p≤0.001, **** p≤0.0001) n=9. **(B)** One way ANOVA indicated this treatment did not affect A6 cell viability as determined by MTT assay, n=12. **(C)** To further confirm, mBCI was used to visualise glutathione content in A6 cells. Representative images at 20x.

**Figure A4:**
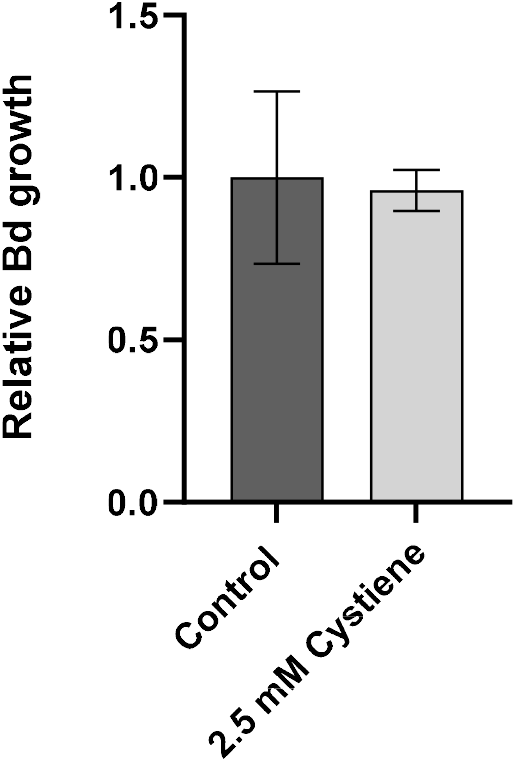
Glutathione supplementation of infected cells does not reduce growth. A6 cells were infected with *Bd* zoospores and infected allowed to progress for 24 hours, after which cells were supplemented with 2.5 mM cysteine for 3 days. *Bd* growth was quantified by staining with calcofluor white and the relative area of stained *Bd* measured using image J from the average of two images per well of three replicate wells, n=6.

## Notes

### Competing Interest Statement

The authors have declared no competing interest.

## References

1. Ku JWK, Gan YH. New roles for glutathione: Modulators of bacterial virulence and pathogenesis. Redox Biol. 2021;44:102012.

2. Wangsanut T, Pongpom M. The Role of the Glutathione System in Stress Adaptation, Morphogenesis and Virulence of Pathogenic Fungi. Int J Mol Sci. 2022;23(18).

3. Thomas D, Klein K, Manavathu E, Dimmock JR, Mutus B. Glutathione levels during thermal induction of the yeast-to-mycelial transition in Candida albicans. FEMS Microbiology Letters. 1991;77(2):331–4.

4. Manavathu M, Manavathu E, Gunasekaran S, Porte Q, Gunasekaran M. Changes in glutathione metabolic enzymes during yeast-to-mycelium conversion of Candida albicans. Canadian Journal of Microbiology. 1996;42(1):76–9.

5. Pócsi I, Prade RA, Penninckx MJ. Glutathione, altruistic metabolite in fungi. Adv Microb Physiol. 2004;49:1–76.

6. Sato I, Shimizu M, Hoshino T, Takaya N. The glutathione system of Aspergillus nidulans involves a fungus-specific glutathione S-transferase. J Biol Chem. 2009;284(12):8042–53.

7. Lee J, Dawes IW, Roe JH. Isolation, expression, and regulation of the pgr1(+) gene encoding glutathione reductase absolutely required for the growth of Schizosaccharomyces pombe. J Biol Chem. 1997;272(37):23042–9.

8. Thangamani S, Eldesouky HE, Mohammad H, Pascuzzi PE, Avramova L, Hazbun TR, et al. Ebselen exerts antifungal activity by regulating glutathione (GSH) and reactive oxygen species (ROS) production in fungal cells. Biochimica et Biophysica Acta (BBA) - General Subjects. 2017;1861(1, Part A):3002–10.

9. Herzenberg LA, De Rosa SC, Dubs JG, Roederer M, Anderson MT, Ela SW, et al. Glutathione deficiency is associated with impaired survival in HIV disease. Proc Natl Acad Sci U S A. 1997;94(5):1967–72.

10. Morris D, Khurasany M, Nguyen T, Kim J, Guilford F, Mehta R, et al. Glutathione and infection. Biochim Biophys Acta. 2013;1830(5):3329–49.

11. Kim VY, Batty A, Li J, Kirk SG, Crowell SA, Jin Y, et al. Glutathione Reductase Promotes Fungal Clearance and Suppresses Inflammation during Systemic Candida albicans Infection in Mice. The Journal of Immunology. 2019;203(8):2239–51.

12. Hiruma K, Fukunaga S, Bednarek P, Piślewska-Bednarek M, Watanabe S, Narusaka Y, et al. Glutathione and tryptophan metabolism are required for *Arabidopsis* immunity during the hypersensitive response to hemibiotrophs. PNAS. 2013;110(23):9589–94.

13. Scheele BC, Pasmans F, Skerratt LF, Berger L, Martel A, Beukema W, et al. Amphibian fungal panzootic causes catastrophic and ongoing loss of biodiversity. Science. 2019;363(6434):1459–63.

14. Zipkin EF, DiRenzo GV, Ray JM, Rossman S, Lips KR. Tropical snake diversity collapses after widespread amphibian loss. Science. 2020;367(6479):814–6.

15. Springborn MR, Weill JA, Lips KR, Ibáñez R, Ghosh A. Amphibian collapses increased malaria incidence in Central America*. Environmental Research Letters. 2022;17(10):104012.

16. Berger L, Skerratt LF, Kosch TA, Brannelly LA, Webb RJ, Waddle AW. Advances in Managing Chytridiomycosis for Australian Frogs: Gradarius Firmus Victoria. Annual Review of Animal Biosciences. 2024;12(1):113–33.

17. Robinson KA, Prostak SM, Campbell Grant EH, Fritz-Laylin LK. Amphibian mucus triggers a developmental transition in the frog-killing chytrid fungus. Current Biology. 2022;32(12):2765-71.e4.

18. Ellison AR, DiRenzo GV, McDonald CA, Lips KR, Zamudio KR. First in Vivo Batrachochytrium dendrobatidis Transcriptomes Reveal Mechanisms of Host Exploitation, Host-Specific Gene Expression, and Expressed Genotype Shifts. G3 (Bethesda). 2017;7(1):269–78.

19. Farrer RA, Martel A, Verbrugghe E, Abouelleil A, Ducatelle R, Longcore JE, et al. Genomic innovations linked to infection strategies across emerging pathogenic chytrid fungi. Nat Commun. 2017;8(1):14742.

20. Webb RJ, Rush C, Berger L, Skerratt LF, Roberts AA. Glutathione is required for growth and cadmium tolerance in the amphibian chytrid fungus, Batrachochytrium dendrobatidis. Biochimie. 2023;220:22–30.

21. Yang X, Wang Y, Zhang Y, Lee W-H, Zhang Y. Rich diversity and potency of skin antioxidant peptides revealed a novel molecular basis for high-altitude adaptation of amphibians. Sci Rep. 2016;6(1):19866.

22. Ren T, Zhu H, Tian L, Yu Q, Li M. Candida albicans infection disturbs the redox homeostasis system and induces reactive oxygen species accumulation for epithelial cell death. FEMS Yeast Research. 2019;20(4).

23. Humphries JE, Melvin SD, Lanctôt C, McCallum H, Newell D, Grogan LF. Chytridiomycosis disrupts metabolic responses in amphibians at metamorphic climax. Microbes and Infection. 2025;27(3):105438.

24. Prostak SM, Fritz-Laylin LK. Laboratory Maintenance of the Chytrid Fungus Batrachochytrium dendrobatidis. Curr Protoc. 2021;1(12):e309.

25. Webb RJ, Vu AL, Skerratt LF, Berger L, De Jesús Andino F, Robert J. Stable in vitro fluorescence for enhanced live imaging of infection models for Batrachochytrium dendrobatidis. PLoS One. 2024;19(8):e0309192.

26. Sumanasekera KK, Skerratt LF, Berger L, Webb RJ. Evaluation of in-vitro growth assays for Batrachochytrium dendrobatidis revealed methylene blue colorimetry is the most accurate. J Microbiol Methods. 2026;241:107394.

27. Webb RJ, Cuf C, Berger L. Glutathione-Mediated Metal Tolerance in an Amphibian Chytrid Fungus (Batrachochytrium dendrobatidis). Environmental Toxicology and Chemistry. 2024;43(7):1583–91.

28. Verbrugghe E, Van Rooij P, Favoreel H, Martel A, Pasmans F. In vitro modeling of Batrachochytrium dendrobatidis infection of the amphibian skin. PLOS ONE. 2019;14(11):e0225224.

29. Verbrugghe E, Pasmans F, Martel A. In Vitro Infection Model Using A6 Cells Sets the Stage for Host-Batrachochytrium salamandrivorans Exploration. J Fungi (Basel). 2025;11(2).

30. Shrieve DC, Bump EA, Rice GC. Heterogeneity of cellular glutathione among cells derived from a murine fibrosarcoma or a human renal cell carcinoma detected by flow cytometric analysis. J Biol Chem. 1988;263(28):14107–14.

31. Neibert KD, and Maysinger D. Mechanisms of cellular adaptation to quantum dots – the role of glutathione and transcription factor EB. Nanotoxicology. 2012;6(3):249–62.

32. Kim H, Xue X. Detection of Total Reactive Oxygen Species in Adherent Cells by 2’,7’-Dichlorodihydrofluorescein Diacetate Staining. J Vis Exp. 2020(160).

33. Shirrif CS, Heikkila JJ. Characterization of cadmium chloride-induced BiP accumulation in Xenopus laevis A6 kidney epithelial cells. Comparative Biochemistry and Physiology Part C: Toxicology & Pharmacology. 2017;191:117–28.

34. Yildiz D, Arik M, Cakir Y, Civi Z. Comparison of N-acetyl-L-cysteine and L-cysteine in respect to their transmembrane fluxes. Biochemistry (Moscow) Supplement Series A: Membrane and Cell Biology. 2009;3(2):157–62.

35. Wang ST, Chen HW, Sheen LY, Lii CK. Methionine and cysteine afect glutathione level, glutathione-related enzyme activities and the expression of glutathione S-transferase isozymes in rat hepatocytes. J Nutr. 1997;127(11):2135–41.

36. Sumanasekera KK, Berger L, Vu AL, Robert J, Akram N, Skerratt LF, et al. Using fluorescent in vitro amphibian cell infection models to quantify Batrachochytrium dendrobatidis pathogenicity. Methods. 2026.

37. De Jesús Andino F, Davydenko A, Webb RJ, Robert J. The Binding, Infection, and Promoted Growth of Batrachochytrium dendrobatidis by the Ranavirus FV3. Viruses. 2024;16(1).

38. Fisher MC, Bosch J, Yin Z, Stead DA, Walker J, Selway L, et al. Proteomic and phenotypic profiling of the amphibian pathogen Batrachochytrium dendrobatidis shows that genotype is linked to virulence. Mol Ecol. 2009;18(3):415–29.

39. Voyles J. Phenotypic profiling of Batrachochytrium dendrobatidis, a lethal fungal pathogen of amphibians. Fungal Ecology. 2011;4(3):196–200.

40. Langhammer PF, Lips KR, Burrowes PA, Tunstall T, Palmer CM, Collins JP. A Fungal Pathogen of Amphibians, Batrachochytrium dendrobatidis, Attenuates in Pathogenicity with In Vitro Passages. PLOS ONE. 2013;8(10):e77630.

41. Davidson MJ, Kosch TA, Aquilina A, Webb RJ, Skerratt LF, Berger L. Influence of Batrachochytrium dendrobatidis isolate and dose on infection outcomes in a critically endangered Australian amphibian. Fungal Ecology. 2025;73:101397.

42. Greener MS, Verbrugghe E, Kelly M, Blooi M, Beukema W, Canessa S, et al. Presence of low virulence chytrid fungi could protect European amphibians from more deadly strains. Nat Commun. 2020;11(1):5393.

43. Wong J, Chen Y, Gan Y-H. Host Cytosolic Glutathione Sensing by a Membrane Histidine Kinase Activates the Type VI Secretion System in an Intracellular Bacterium. Cell Host & Microbe. 2015;18(1):38–48.

44. Fratelli M, Demol H, Puype M, Casagrande S, Eberini I, Salmona M, et al. Identification by redox proteomics of glutathionylated proteins in oxidatively stressed human T lymphocytes. Proc Natl Acad Sci U S A. 2002;99(6):3505–10.

45. Claytor S. The role of serotonin and glutathione in the pathogenesis of chytridiomycosis. PhD thesis. 2020.

46. Seefeldt T, Zhao Y, Chen W, Raza AS, Carlson L, Herman J, et al. Characterization of a novel dithiocarbamate glutathione reductase inhibitor and its use as a tool to modulate intracellular glutathione. J Biol Chem. 2009;284(5):2729–37.

47. Gansemer ER, McCommis KS, Martino M, King-McAlpin AQ, Potthof MJ, Finck BN, et al. NADPH and Glutathione Redox Link TCA Cycle Activity to Endoplasmic Reticulum Homeostasis. iScience. 2020;23(5):101116.

48. Muller E. A glutathione reductase mutant of yeast accumulates high levels of oxidized glutathione and requires thioredoxin for growth. Molecular biology of the cell. 1996;7(11):1805–13.

49. Black B, da Silva LBR, Hu G, Qu X, Smith DFQ, Magaña AA, et al. Glutathione-mediated redox regulation in Cryptococcus neoformans impacts virulence. Nature Microbiology. 2024;9(8):2084–98.

50. Wang X, Che MZ, Khalil HB, McCallum BD, Bakkeren G, Rampitsch C, et al. The role of reactive oxygen species in the virulence of wheat leaf rust fungus Puccinia triticina. Environ Microbiol. 2020;22(7):2956–67.

51. Zhang Z, Chen Y, Li B, Chen T, Tian S. Reactive oxygen species: A generalist in regulating development and pathogenicity of phytopathogenic fungi. Computational and Structural Biotechnology Journal. 2020;18:3344–9.

52. James MR, Doss KE, Cramer RA. New developments in Aspergillus fumigatus and host reactive oxygen species responses. Current Opinion in Microbiology. 2024;80:102521.

53. Ding Y, Yan B, Zhao S, Chen Y, Wan H, Qian W. Synthetic modulation of ROS scavenging during host— Sclerotinia sclerotiorum interaction: a new strategy for the development of highly resistant plants. Phytopathology Research. 2024;6(1):20.

54. McDonald CA, Ellison AR, Toledo LF, James TY, Zamudio KR. Gene expression varies within and between enzootic and epizootic lineages of Batrachochytrium dendrobatidis (Bd) in the Americas. Fungal Biology. 2020;124(1):34–43.

55. Bolter C, Brammall RA, Cohen R, Lazarovits G. Glutathione alterations in melon and tomato roots following treatment with chemicals which induce disease resistance to Fusarium wilt. Physiological and Molecular Plant Pathology. 1993;42(5):321–36.

56. McMahon TA, Halstead NT, Johnson S, Rafel TR, Romansic JM, Crumrine PW, et al. The Fungicide Chlorothalonil Is Nonlinearly Associated with Corticosterone Levels, Immunity, and Mortality in Amphibians. Environmental Health Perspectives. 2011;119(8):1098–103.

57. Jiménez RR, Alvarado G, Ruepert C, Ballestero E, Sommer S. The Fungicide Chlorothalonil Changes the Amphibian Skin Microbiome: A Potential Factor Disrupting a Host Disease-Protective Trait. Applied Microbiology. 2021;1(1):26–37.

58. Van Meter RJ, Glinski DA, Purucker ST, Henderson WM. Induced Hepatic Glutathione and Metabolomic Alterations Following Mixed Pesticide and Fertilizer Exposures in Juvenile Leopard Frogs (Lithobates sphenocephala). Environ Toxicol Chem. 2022;41(1):122–33.

59. Prokić MD, Borković-Mitić SS, Krizmanić II, Mutić JJ, Vukojević V, Nasia M, et al. Antioxidative responses of the tissues of two wild populations of Pelophylax kl. esculentus frogs to heavy metal pollution. Ecotoxicology and Environmental Safety. 2016;128:21–9.

60. Esmaeilbeigi M, R PD, B JK, Ezaz T, Clulow S. Evidence for a metal disease refuge: The amphibian-killing fungus (Batrachochytrium dendrobatidis) is inhibited by environmentally-relevant concentrations of metals tolerated by amphibians. Environ Res. 2024;261:119752.

